# Lung organoids and microplastic fibers: a new exposure model for emerging contaminants

**DOI:** 10.1101/2021.03.07.434247

**Authors:** Anna Winkler, Nadia Santo, Laura Madaschi, Alessandro Cherubini, Francesco Rusconi, Lorenzo Rosso, Paolo Tremolada, Lorenza Lazzari, Renato Bacchetta

**Author notes:** co-last author for the human organoids. co-last-author for microplastic and microscopy analyses. Address for correspondence to Anna Winkler, Department of Environmental Science and Policy, University of Milan, Via Celoria 26, 20133 Milan, Italy.

## Abstract

**Background:** Three-dimensional (3D) structured organoids are the most advanced *in vitro* models for studying human health effects, but they have been applied only once to evaluate the biological effects associated with microplastic exposure. Fibers from synthetic clothes and fabrics are a major source of airborne microplastics, and their release from dryer machines is still poorly understood.

**Objectives:** In this study, we aimed to establish an *in vitro* organoid model of human lung epithelial cells to evaluate its suitability for studying the effects of airborne microplastic contamination on humans. Furthermore, we aimed to characterize the microplastic fibers (MPFs) released in the exhaust filter of a household dryer and to test their interactions and inflammatory effects on the established lung organoids.

**Methods:** The polyester fibers emitted from the drying of synthetic fabrics were collected. Morphological characterization of the fibers released into the air filter was performed by optical microscopy and scanning electron microscopy (SEM)/energy dispersive x-ray spectroscopy (EDS). The organoids were exposed to various MPF concentrations (1, 10, and 50 mg L^−1^) and analyzed by optical microscopy, SEM, and confocal microscopy. Gene expression analysis of lung-specific genes, inflammatory cytokines, and oxidative stress-related genes was achieved by quantitative reverse transcription–polymerase chain reaction (qRT-PCR).

**Results:** We successfully cultured organoids with lung-specific genes. The presence of MPFs did not inhibit organoid growth, but polarized cell growth was observed along the fibers. Moreover, the MPFs did not cause inflammation or oxidative stress. Interestingly, the MPFs were coated with a cellular layer, resulting in the inclusion of fibers in the organoid.

**Discussion:** This work could have potential long-term implications regarding lung epithelial cells undergoing repair. This preliminary exposure study using human lung organoids could form the basis for further research regarding the toxicological assessment of emerging contaminants such as micro- or nanoplastics.

## Introduction

Atmospheric contamination through airborne particles and fibers is a long-existing and growing research topic of environmental pollution. Several epidemiological studies have already linked air pollution through ambient atmospheric particles to many adverse human health effects, including respiratory illness, cardiovascular disease, and carcinogenic effects (Churg and Bauer 2000; Loomis et al. 2013; Valavandis et al. 2008). In addition to the well-known air pollutants such as combustion particles from fuel-burning emissions and other particulate organic matter and aerosols (Nel 2005), microplastics (MPs) are now considered as emerging components of air pollution (Zhang et al., 2020 and authors therein).

The human health effects of MPs and nanoplastics (NPs) are usually deduced from *in vivo* exposure tests in animal models, i.e., mammalian or nonmammalian models such as mice, rats, *Xenopus laevis* (De Felice et al. 2018), and zebrafish (Bhagat et al. 2020), and from *in vitro* models such as human cell cultures. After exposure, i.e., inhalation or ingestion, MPs/NPs may pass through biological barriers, leading to translocation in the body tissue. Indeed, the transfer of MPs/NPs into human cells has been demonstrated using a variety of human cell lines, including epidermal cells (Triebskorn et al. 2019), lung epithelial cells (Lim et al. 2019; Schirinzi et al. 2017; Xu et al. 2019), endothelial cells (Barshtein et al. 2016), and intestinal cells (Cortés et al. 2020; Domenech et al. 2020). Human two-dimensional (2D) cell cultures are common *in vitro* models used to evaluate the biological effects associated with MP/NP exposure; however, these systems have some limitations such as an inaccurate representation of the *in vivo* tissue (Costa et al. 2016). Very recently, organoids, which are three-dimensional (3D) cellular structures generated from induced pluripotent stem cells, embryonic stem cells, or adult tissue-resident stem cells (Shamir and Ewald 2014), have been shown to be a powerful tool to overcome the limits of 2D culture. Indeed, organoids, such as kidney (Takasato et al. 2016), brain (Lancaster et al. 2013), intestine (Sato et al. 2011), liver (Huch et al. 2015), pancreas (Dossena et al. 2020), and lung (Sachs et al. 2019) organoids, have recently emerged as attractive model systems that contain key aspects of *in vivo* tissue and organ complexity while being more experimentally manageable than model organisms. Moreover, these 3D structures have been recently applied in nanosafety research (Kämpfer et al. 2020) and modeling diseases such as cancer (Tuveson and Clevers 2019). Nevertheless, organoids have been applied only once to evaluate the biological effects associated with MP/NP exposure. The recent study by van Dijk et al. (2021) tested the effects of nylon and polyester fibres on murine and human lung organoids showing inhibited organoid growth (lower organoid number and smaller size, evaluated by light and fluorescence microscopy observations). For nylon, the effects were more pronounced due to component leaching of the polymer but were limited to organoids in development phase. The study applied gene expression pathway analysis of organoids exposed to nylon fibers to confirm the results.

MPs are ubiquitous in the environment (Prata et al. 2020). In the atmosphere, a large proportion of the MPs consists of microplastic fibers (MPFs) derived from various sources, including synthetic clothes, textiles, and upholstery (Henry et al. 2019). Undeniably, synthetic clothing mostly made of polyethylene terephthalate (namely, polyester) is considered to be a major source of airborne MPs (Dris et al., 2016). Recent studies also have demonstrated that a major pathway for MPFs is the atmosphere, through which they can reach very remote areas such as glaciers (Ambrosini et al. 2019), the Arctic region (Bergmann et al., 2017; Lusher et al., 2015), and Antarctica (Reed et al., 2018; Waller et al., 2017). Furthermore, Dris et al. (2017) and Liu et al. (2019) have investigated and compared MPFs in indoor and outdoor air. They found that indoor air contains considerably higher amounts of natural and synthetic fibers than outdoor air and dust.

MPFs can be released into the wastewater through the washing of synthetic textiles and clothes, as demonstrated previously in several studies (Browne et al., 2011; De Falco et al., 2019; among others). In addition to the washing process, the drying of synthetic textiles with a household clothes dryer further emits MPFs into the atmosphere via the exhaust air. However, the air contamination caused by the drying of synthetic clothes and fabrics is still poorly understood. Recently, O’Brien et al. (2020) have reported and quantified for the first time the amount of MPFs released from a clothes dryer into the ambient air. The issue was further addressed by Kapp and Miller (2020), who studied the spatial distribution of MPFs emitted from the vent of a clothes dryer directly into the environment.

Human inhalation of atmospheric MPFs has been demonstrated *in vivo* (human lung biopsies) by Pauly et al. (1998) and Pimentel et al. (1975). The deposition of MPs, however, depends on the properties and lung anatomy (Churg and Bauer 2000; Lippmann et al. 1980). Currently, it is well known that humans in certain exposure scenarios, such as industrial workers, are particularly susceptible to diseases caused by airborne MPs in the lung (Eschenbacher et al. 1999; Hours et al. 2007). In the attempt to estimate the human inhalation of indoor airborne MPs, Vianello et al. (2019) set up a mannequin simulating the human metabolic rate and breathing (male person with light activity). The simulated human exposure revealed an average inhaled concentration of 9.3 ± 5.8 MPs m^−3^ (or 272 MPs per day), corresponding to the value of indoor airborne fibers reported by Dris et al., 2017 (median value of 5.4 fibers m^−3^) - disregarding the problem of comparing the results of studies using different analytical techniques.

The available data or information providing evidence of the negative human health effects of inhaled MPFs are still rare and insufficient (Prata 2018). To date, the human exposure risk still remains unclear, and the consequences of MPF exposure are not yet well understood (Prata et al. 2020). Accordingly, there is an increasing demand for interdisciplinary research between environmental and human health sciences (Dries et al. 2017). In this study, we propose the application of the innovative human organoid model for human exposure tests for MPs/NPs and other particulate contaminants. Considering the described exposure risk for humans to airborne MPs, the use of the human lung as an organoid model in this study is an innovative experimental approach.

Importantly, exposure tests should not only apply environmentally relevant concentrations but also environmentally relevant MP properties (Prata et al. 2020), such as nonstandard, fibrous MPs. Therefore, to test the effects of airborne MPFs on human lung airway organoids (hAOs), polyester fibers emitted from the drying process of synthetic clothes and fabrics were collected and used as a test contaminant. The specific aims of this paper were as follows: i) to characterize the release of MPFs in the exhaust filter of a household dryer machine; ii) to analyze the effect of these environmentally relevant MPFs on hAOs as a possible target of airborne contamination from synthetic clothes; iii) to characterize the established hAOs; and iv) to evaluate the use of human organoid models to assess the effects of pollutants to humans.

## Material and methods

### hAO isolation and culture

The herein applied hAOs were generated from tissue-resident adult stem cells. Healthy human lung tissues (airway space) were obtained from the Thoracic Surgery and Lung Transplant Unit, IRCCS Ca’ Granda Ospedale Maggiore Policlinico, Milan, Italy. The use of human specimens was approved by the Institutional Review Board (CE 0001977). Young patients underwent minimal invasive wedge lung resection for spontaneous pneumothorax.

For the processing of solid lung tissue, the biopsies (0.5–1 cm^3^) were minced in small pieces and rinsed with wash medium (Dulbecco’s modified Eagle medium, DMEM) supplemented with 1% fetal bovine serum, 1% penicillin/streptomycin and 1× Glutamax (see Table S1 for the source and ID numbers of all reagents used). Fragments were digested in wash medium containing 0.125 mg mL^−1^ collagenase I, 0.125 mg mL^−1^ dispase II, and 0.1 mg mL^−1^ DNase I for 90 min at 37 °C. The digestion was stopped by adding cold wash medium, and the suspension was filtered through a 70-μm cell strainer and then spun for 5 min at 400 × *g.* In case of visible red pellets, erythrocytes were lysed in 5 mL of red blood cell lysis buffer for 15 min at room temperature before the addition of wash medium and centrifugation at 400 × *g.* The cell pellet was mixed with Matrigel, and 40 μL of Matrigel-cell suspension was allowed to solidify on prewarmed nontissue culture 24-well plates for 20-30 min. After complete Matrigel solidification, culture medium containing AdDMEM/F12 supplemented with 10 mM Hepes, 1× penicillin/streptomycin, 1× Glutamax, 1% B27, 1.25 mM N-acetylcysteine, 25 ng mL^−1^ FGF7, 500 ng mL^−1^ RSPO1, 100 ng mL^−1^ FGF10, 100 ng mL^−1^ Noggin, 5 mM nicotinamide, 500 nM A83.01, and 500 nM SB202190 was added. For establishment of the organoids, the culture medium was supplemented with 10 μM Rock inhibitor Y27632 during the first three days. The culture medium was changed every 3-4 days. After 10-14 days, the organoids were removed from the Matrigel, mechanically dissociated into small fragments, and then split 1:4-1:6 in fresh Matrigel, enabling the formation of new organoids. All cell cultures were routinely tested for mycoplasma contamination by quantitative reverse transcription-polymerase chain reaction (qRT-PCR).

### hAO characterization

Total RNA from organoids was isolated using TRIzol reagent, according to the manufacturer’s instructions. The RNA concentration and purity were verified using a NanoDrop ND-100 spectrophotometer (NanoDrop Technologies). For the qRT-PCR assay, cDNA was synthesised from 200 ng of total RNA with SuperScript IV VILO. The cDNA was diluted 10-fold, and 1 μL of the sample was used as a template for qRT-PCR analysis with SYBR Select Master Mix on a CFX96 thermal cycler (Bio-Rad). The relative expression levels of the selected target genes were determined using the ΔΔC_t_ method and normalized to the geometric mean of the ACTB and TBP mRNA levels using the primers listed in Table S2.

To characterize the hAO, gene expression analysis by qRT-PCR was performed after 10-14 days of standard culture and again after MPF exposure to investigate possible changes in the lung cell compartments.

### Sampling and characterization of MPFs from a dryer machine

To generate environmentally relevant MPFs for the human organoid exposure experiment, polyester fibers emitted from the drying of synthetic clothes and fabrics were collected. Specifically, one polyester t-shirt, six sweatshirts, and two blankets of different colors (dry weight: 5,427 g) were washed in a washing machine and subsequently dried in a common domestic tumble dryer. The dryer filter was previously cleaned by a vacuum cleaner, and all fibers derived from the first drying cycle were discarded. The same items were then washed again, and the synthetic fibers from the filtered exhaust air of the tumble dryer were collected (Figure S1), wrapped in aluminum foil, and transported to the laboratory for subsequent analyses.

The MPFs were first morphologically analyzed with a Leica EZ4D stereomicroscope and then with a Leica DMRA2 light microscope equipped with a Leica DC300 F digital camera. The detailed morphology and the elemental composition of the fibers were studied with a Zeiss LEO 1430 scanning electron microscope (SEM) coupled with a Centaurus detector for energy dispersive x-ray spectroscopy (EDS) analysis. A subsample of MPFs was mounted onto standard SEM stubs and gold-coated. The elemental analysis was performed using Oxford Instruments INCA version 4.04 software (Abingdon, UK). The operating conditions were as follows: accelerating voltage, 20 kV; probe current, 360 pA; and working distance, 15.0 mm.

### MPF exposure to hAOs

The MPFs collected from the filter of the dryer machine were resuspended at a final concentration of 500 mg L^−1^ in AdDMEM/F12 supplemented with 10 mM Hepes, 1× penicillin/streptomycin, and 1× Glutamax; then the sample was sonicated at a high intensity (3 × 5 min) using a Bioruptor sonicator (Diagenode). To obtain organoid-MPF cocultures, the hAOs were split as described above, and then the organoid fragments were embedded in Matrigel drops and mixed with MPFs at a ratio of 70% Matrigel with organoids and 30% culture medium containing the MPFs, at different concentrations (1, 10, and 50 mg L^−1^). Upon solidification of the Matrigel, the organoid-MPF coculture was supplemented again with the culture medium containing the MPFs at the respective concentrations. After ten days, the organoids reached maturation (ca. 200-300 μm in diameter). At this point, in order to improve the interaction between the organoids and MPFs, the hAOs were carefully removed from the Matrigel and cultured in suspension with only culture medium containing the MPFs at the same concentrations for 1 week with gentle agitation. An organoid coculture was cultivated under the same conditions without MPF exposure as a control. After exposure, the lung organoid-MPF cocultures were collected for gene expression analysis and fixed for confocal microscopy and SEM analysis, as described below.

### SEM

To study the effects of MPFs on organoid growth, the control and exposed samples (1, 10, and 50 mg L^−1^) were fixed in a mixture of 4% paraformaldehyde and 2% glutaraldehyde in 0.1 M sodium cacodylate-buffered solution at pH 7.4. After several washes in the same buffer, the samples were post-fixed in 1% OsO_4_ for 1.5 h at 4°C and then dehydrated in a graded ethanol series. As a final step, the organoids were treated with hexamethyldisilazane for complete chemical dehydration. All samples were mounted onto standard aluminum stubs, gold sputtered, and analyzed under a Zeiss LEO 1430 SEM at 20 kV.

### Immunofluorescence staining and confocal microscopy

For confocal microscopy analysis, the organoids were washed with 0.01 M phosphate-buffered saline, pH 7.4 (PBS) and fixed for 15 min in 4% paraformaldehyde, incubated in 0.05 M NH_4_Cl in PBS for 30 min, permeabilized for 15 min in PBS containing 1% bovine serum albumin (BSA) and 0.2% Triton X-100, and blocked for 30 min in 1% BSA. The organoids were incubated overnight at 4 °C with a mouse monoclonal antibody against α-tubulin diluted in 0.1% BSA/PBS. The samples were then rinsed in PBS and incubated for 3 h at room temperature with rhodamine phalloidin (cytoskeleton) and the secondary Alexa Fluor 488- conjugated goat anti-mouse antibody. After several washes with PBS, the organoids were finally incubated for 5 min at room temperature with the DNA dye Hoechst 33342 (1:5000). At the end of the staining procedure, the organoids were washed with PBS (3 × 5 min), mounted in 1:2 PBS/glycerol, and observed under a Nikon A1 confocal microscope.

### Inflammatory cytokine and oxidative stress evaluation

To investigate the possible changes in the inflammatory cytokine expression in hAOs after MPF exposure, gene expression analysis by qRT-PCR was performed as described above at the end of the coculture with the highest MPF concentration using the primers listed in Table S2. Considering that the exposure of MPFs could induce oxidative stress in human tissues (Hu and Palić 2020), we analyzed the expression of genes involved in oxidative stress pathways reported in relation to MPF exposure, including superoxide dismutase family genes (SOD1 and SOD2), glutathione detox-related genes (GSTA1 and GPX1), catalase (CAT), and ROS-controlling genes (NOX2, COX1, and ND1). In addition, the capacity of our 3D structures to respond to standard inflammatory stimuli was evaluated after treatment with poly(I:C) (50 μg mL^−1^) as a positive control.

### Data analysis

For statistical analysis of the gene expression data, a two-way analysis of variance followed by Bonferroni’s multiple comparison test (α = 0.05) was performed. Photoshop was used to export the image graphs.

## Results

### hAO isolation and characterization

The hAOs were successfully generated from the biopsies of three healthy lung donors. A heatmap of selected genes (Figure 1) demonstrated that the hAO lines derived from the three different donors have similar gene expression profiles (compared to human pancreas organoids as a control).

**Figure 1.**
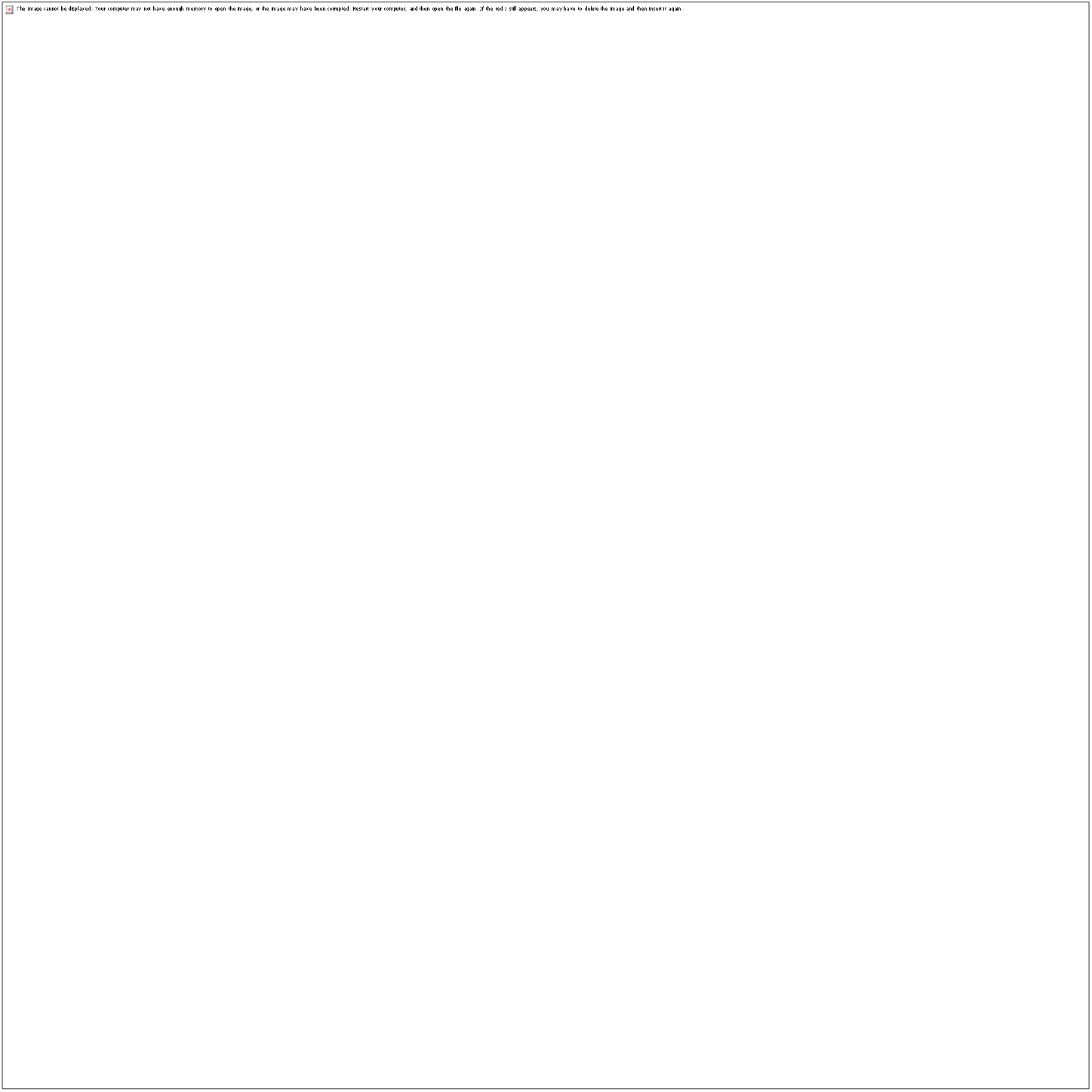
Gene expression heatmap of human lung airway organoids (hAOs) generated from the biopsies of three healthy lung donors and a human pancreas organoid (hPO) for comparison.

The organoids were analyzed by optical microscopy, confocal 3D construction, and SEM imaging. The normal organoids (control group) exhibited a spherical shape with a typical diameter of 200-300 μm and an inner cavity (Figure 2A-B). Confocal microscopy confirmed that the cells were differentiated, showing the presence of ciliated cells and nonciliated cells, including club cells (identified by the presence of microvilli), that were irregularly distributed on the surface and corresponding to their *in vivo* position (Figure 2C-E). Immunofluorescence experiments with the actin cytoskeleton marker revealed the different shapes and dimensions of the cells containing nuclei, which were visible in the surface section (Figure 2C-E) and transverse plane (Figure 2D). Moreover, the inner surface of the organoids established a differentiated cell structure, including ciliated cells facing the organoid cavity, as demonstrated in Figure 2D. The presence of cilia at the inner surface of the organoids could be explained by the rotating motion of the organoid cavity, which can be seen in a video recording of the organoids (Video S1). High-resolution SEM imaging of the organoid surface showed the same irregularly shaped ciliated and nonciliated cells, displaying the dimensions of the cilia and microvilli (Figure 2F).

**Figure 2.**
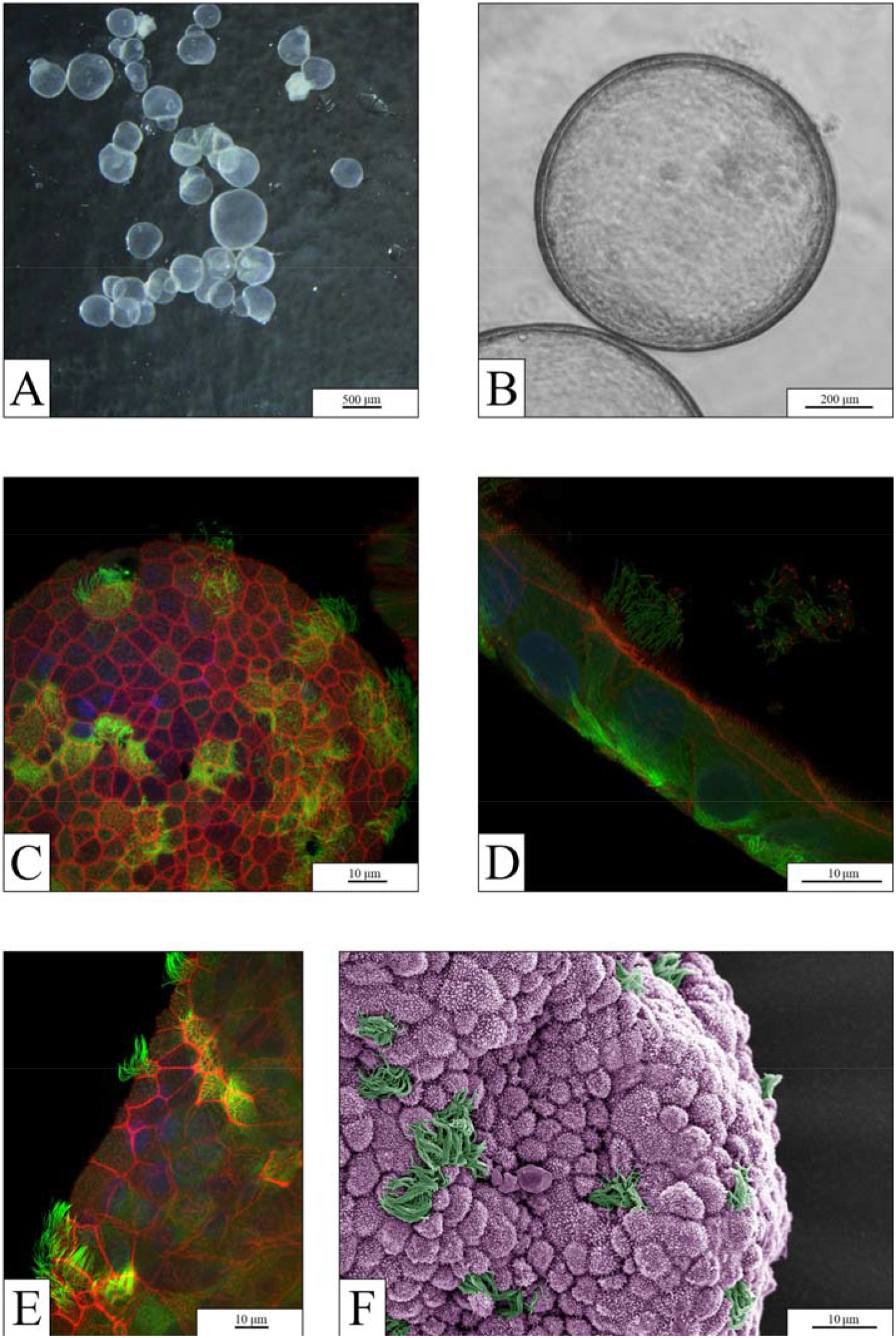
Characterization of human lung organoids (control group). (A) Stereo microscopy image and (B) optical microscopy image of organoids. (C-E) Immunofluorescence imaging of the organoid surface and the transverse section generated by confocal microscopy showing the cellular organization with a cytoskeletal marker (anti-acetylated tubulin; green) and counterstaining of the actin cytoskeleton (phalloidin 565; red) and nuclei (Hoechst 33342; blue). (F) Pseudo-colored SEM image of the organoid’s surface showing irregularly shaped multiciliated (green) and nonciliated cells with microvilli (purple).

### MPFs in the air filter of a dryer machine

The total weight of the MPFs removed from the air filter of a dryer machine was 2.3 g (Figure S1), which was released from 5.4 kg of 100% polyester fabrics (0.42 g of dry MPFs kg^−1^ dry fabric). The procedure was repeated once again with the same fabrics (rewashing and redrying), and the release of MPFs was similarly high. Optical microscopy experiments demonstrated fibers of different colors, reflecting the color mixture of the dried materials (Figure 3A-B). For a more detailed analysis, SEM (Figure 3C-E) revealed that the MPFs varied in size and morphology. The fiber length ranged from 139 μm to 1,535 μm (mean: 663 ± 333 μm). Interestingly, the shape of these MPFs differed at the transverse surface; no MPFs had a round profile along the entire length, but they exhibited a varying profile from flat and twisted to tattered (Figure 3C-E). Flat MPFs exhibited maximum widths of 21-24 μm along their widest dimension and a minimum height of 1-3 μm along their thinnest dimension. Furthermore, we observed that some fibers showed a rough surface, where small pieces of <10 μm flaked off from the fiber (Figure 3D) or a chapped ending was revealed (Figure 3E), which could lead to the release of very small MPs from the fibers.

**Figure 3.**
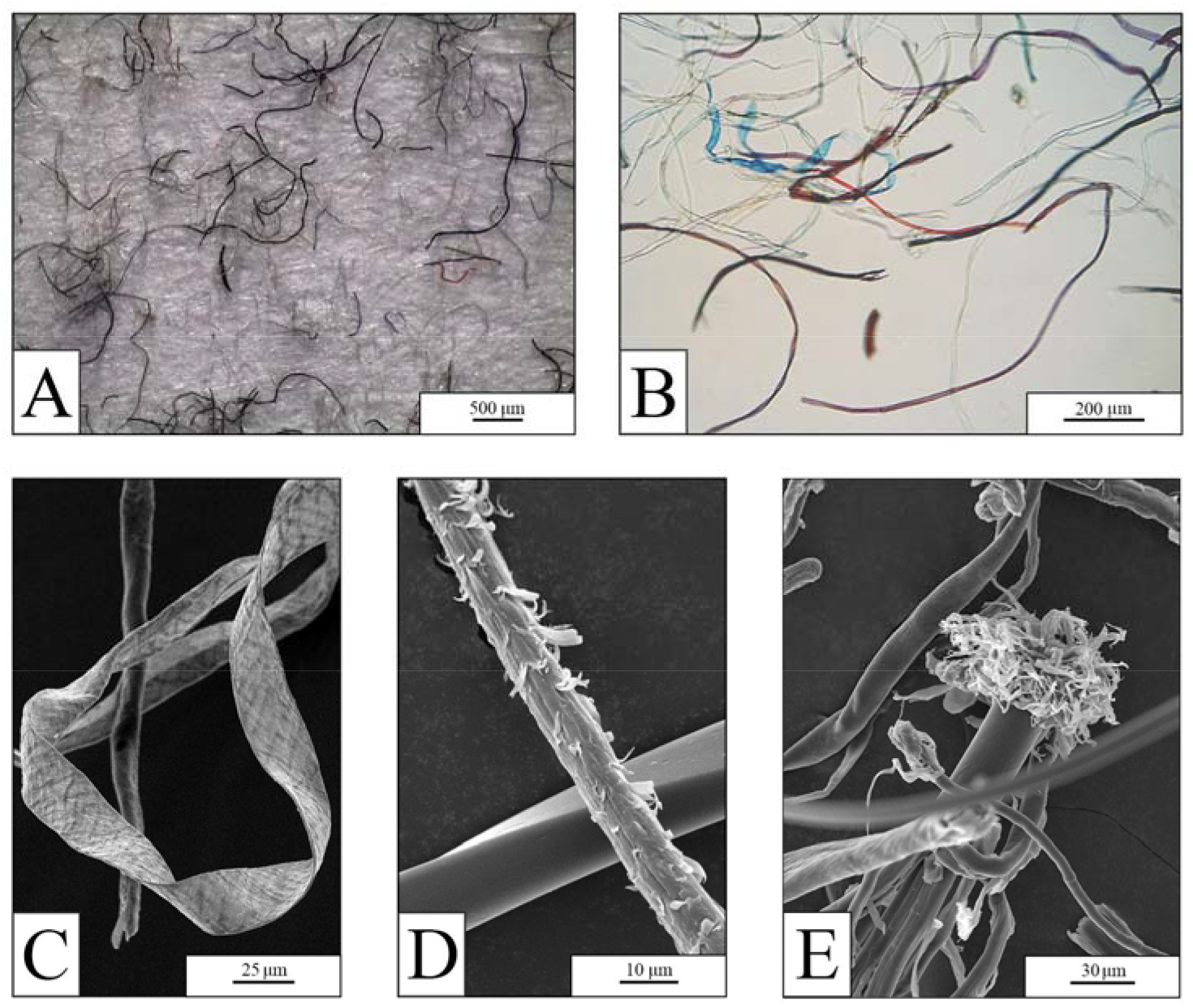
Synthetic fibers obtained from the air filter of a dryer machine. (A-B) Optical microscopy images demonstrating MPFs of different sizes and colors. (C-E) SEM images of MPFs at different magnifications showing varying morphologies.

Considering the average dimensions of the fibers described by SEM analyses, we obtained a fiber volume of 28,485 μm^3^ fiber^−1^ (633 μm × 22.5 μm × 2 μm) or 2.8 × 10^−5^ mm^3^ fiber^−1^, indicating that the 2.3 g of polyester fibers collected in the filter of a dryer machine contained 59 × 10^6^ fibers (considering that the density of polyester is 1.38 mg mm^−3^). Elemental analysis showed a C:O ratio of 74:26, which corresponds to that of polyester (see Figure S2 for the EDS spectrum), thus confirming that the fibers were made of polyester.

### Effects of MPFs on hAOs

The hAOs exposed to MPFs were affected by all concentrations of MPFs (1, 10, and 50 mg L^−1^). Optical microscopy, confocal 3D construction, and SEM image analyses revealed that the organoids exposed to MPFs were not inhibited in their growth and did not exhibit any cellular abnormalities compared to the control group, independent of the MPF concentration (Figure 4A-F). Most organoids did not interact with the fibers, or only to a small extent, and maintained their radial (almost spherical) architecture (Figure 4D-F). In addition, no difference in the expression of epithelial to mesenchymal transition markers was observed (Figure 5A). Moreover, fluorescence staining of the surface (Figure 6A-B) and transverse (Figure 6C) sections of the organoids as well as high-resolution SEM images (Figure 6D-H) showed the same cellular structure as the controls, including ciliated and nonciliated cells, even when the organoids were in contact with MPFs. To verify that the lung heritage of hAOs was not affected by their exposure to MPFs, we performed gene expression analysis of the typical epithelial lung markers NKX2.1 and CLDN1 as well as the specific airway lung markers SFTPA1 and SFTPC (AT2 cells), SCGB1A1 (club cells), NPHP1 and DNAH5 (ciliated cells), and KRT5 (basal cells); no significant differences were observed between the organoids exposed to MPFs and the control organoids (Figure 5B). The only notable observation was a higher density of polarization of the cytoskeleton near the fiber contact site, visible as an intensification of the red color in Figure 6B. Owing to an organoid that had broken apart (likely during the transfer of the dehydrated organoid onto the aluminum stub), we were able to take SEM images of the inner surface of the organoids and could observe the same cellular differentiation (multiciliated and nonciliated cells) as the outer surface (Figure 6E). Rather than depending on the MPF concentration, the observed effects varied with the degree of MPF-organoid contact, which presumably depended on the development phase of the organoids. The organoids that made contact with fibers during maturation grew around the fibers and integrated them into their body (Figure 6F-H).

**Figure 4.**
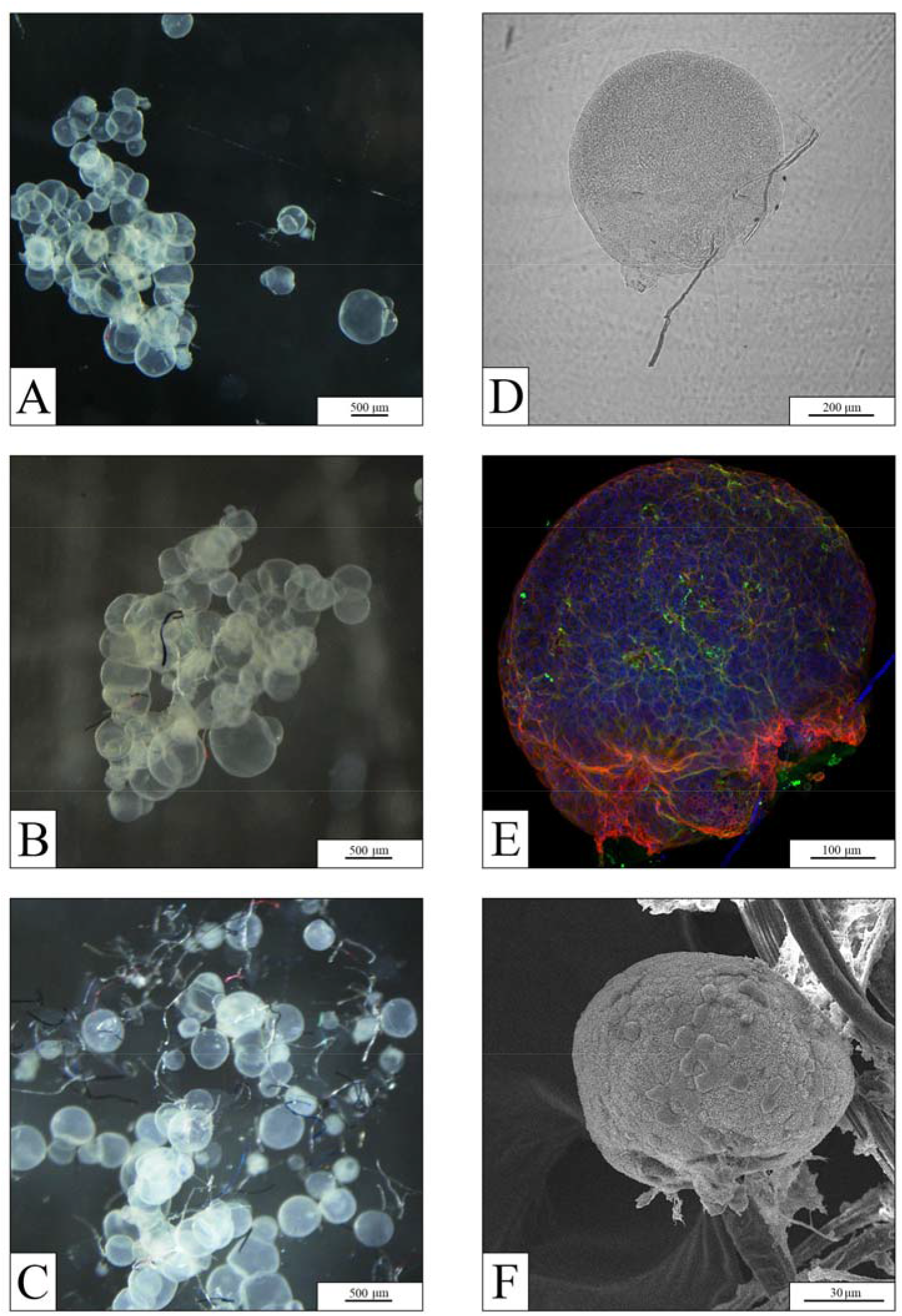
Human lung organoids exposed to MPFs. (A-C) Stereo microscopy images of organoids in suspension with MPFs at increasing concentrations (top to bottom: 1, 10, and 50 mg L^−1^). (D) Phase contrast image and (E) immunolabeled 3D reconstruction of the same organoid (MPF concentration: 10 mg L^−1^) showing the interaction between the organoid and a fiber. The organoid was stained for cytoskeletal proteins: microtubules (anti-acetylated tubulin; green), F-actin (phalloidin 565; red), and nuclei (Hoechst 33342; blue). Also visible in blue is a synthetic fiber. (F) SEM image of an organoid (MPF concentration: 50 mg L^−1^) in contact with a fiber, demonstrating a spherical body and a differentiated cell structure.

**Figure 5.**
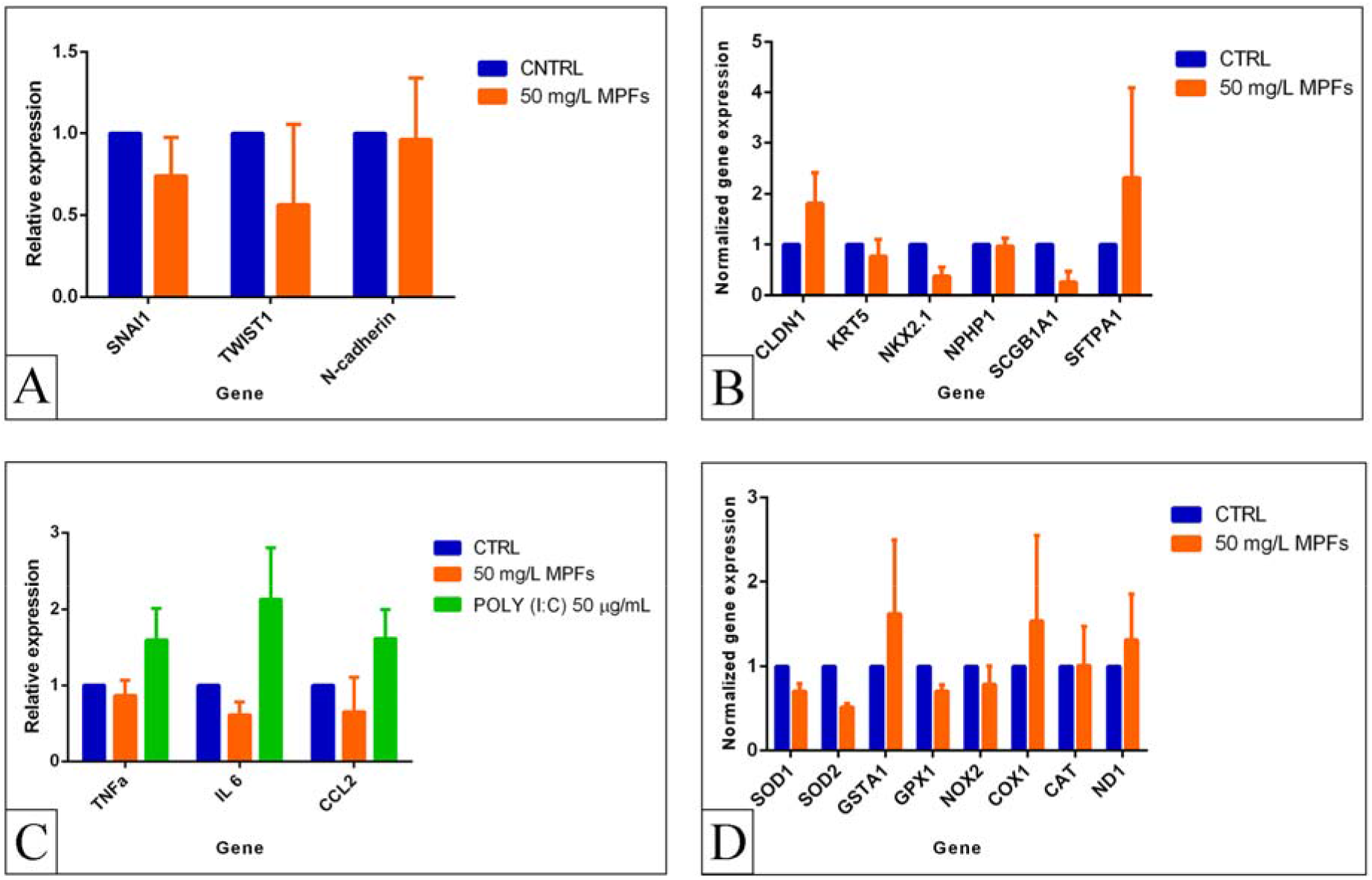
qRT-PCR analysis of the control organoids (blue) compared to organoids exposed to MPFs (red; MPF concentration: 50 mg L^−1^) for selected target genes: (A) epithelial to mesenchymal transition markers, (B) general epithelial lung markers and specific airway markers, (C) inflammatory cytokines compared to the response of positive control organoids to standard inflammatory stimuli (green; poly(I:C) 50 μg mL^−1^), and (D) oxidative stress-related genes.

**Figure 6.**
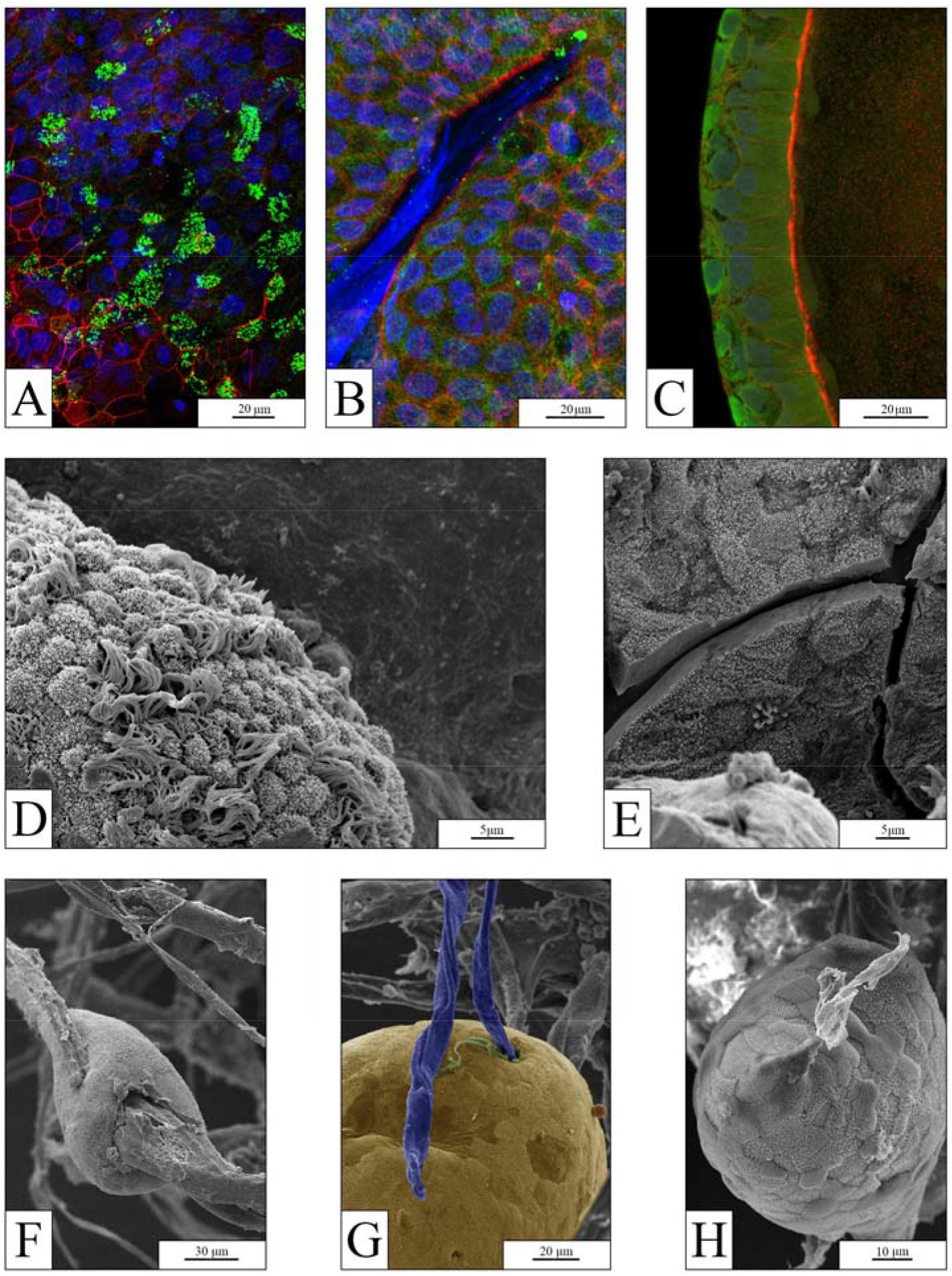
Human lung organoids exposed to MPFs. (A-B) Surface and (C) transverse sections of an immunolabeled organoid (MPF concentration: 50 mg L^−1^) showing the cell organization. The organoid was stained to show ciliated cells (anti-acetylated tubulin; green), F-actin cytoskeleton (phalloidin 565; red), and nuclei (Hoechst 33342; blue). Also visible in blue is a synthetic fiber. (D) SEM image of an organoid’s outer surface showing ciliated and nonciliated cells (MPF concentration: 10 mg L^−1^). (E) SEM image of the inner surface of an organoid that had broken apart, demonstrating cell differentiation facing the organoid’s cavity (MPF concentration: 10 mg L^−1^). (F-H) SEM images of organoids (MPF concentration: 50 mg L^−1^) demonstrating a differentiated cell structure and integration of a fiber into the organoid; the image in panel G is partially pseudo-colored for better visualization (cell tissue, yellow; fiber, blue).

Interestingly, the organoids closely bonded to an MPF formed an organoid-fiber interlacement or fully encompassed the MPF. We further observed that some organoids grew polarized around the fibers. For example, Figure 7A-G shows cell growth along the length of a fiber. The organoids seemingly included and phagocytosed fibers by developing netlike extensions of cells (clearly visible in Figure 7D-F), resulting in a full internalization of the MPFs. While the cells of the organoid body growing along the length of a fiber exhibited a differentiated structure (Figure 7A-B), the thin cell extension covering the surface of a fiber exiting the organoid body (Figure 7D-F) was rather smooth and uniform. These observations of cells covering a fiber with a thin layer stands in contrast to the images depicted in Figure 6F-H showing that cells did not cover the fibers once they left the organoid. We also noticed that organoids in suspension with MPFs can be damaged by fibers piercing the cell tissue (Figure 7C).

**Figure 7.**
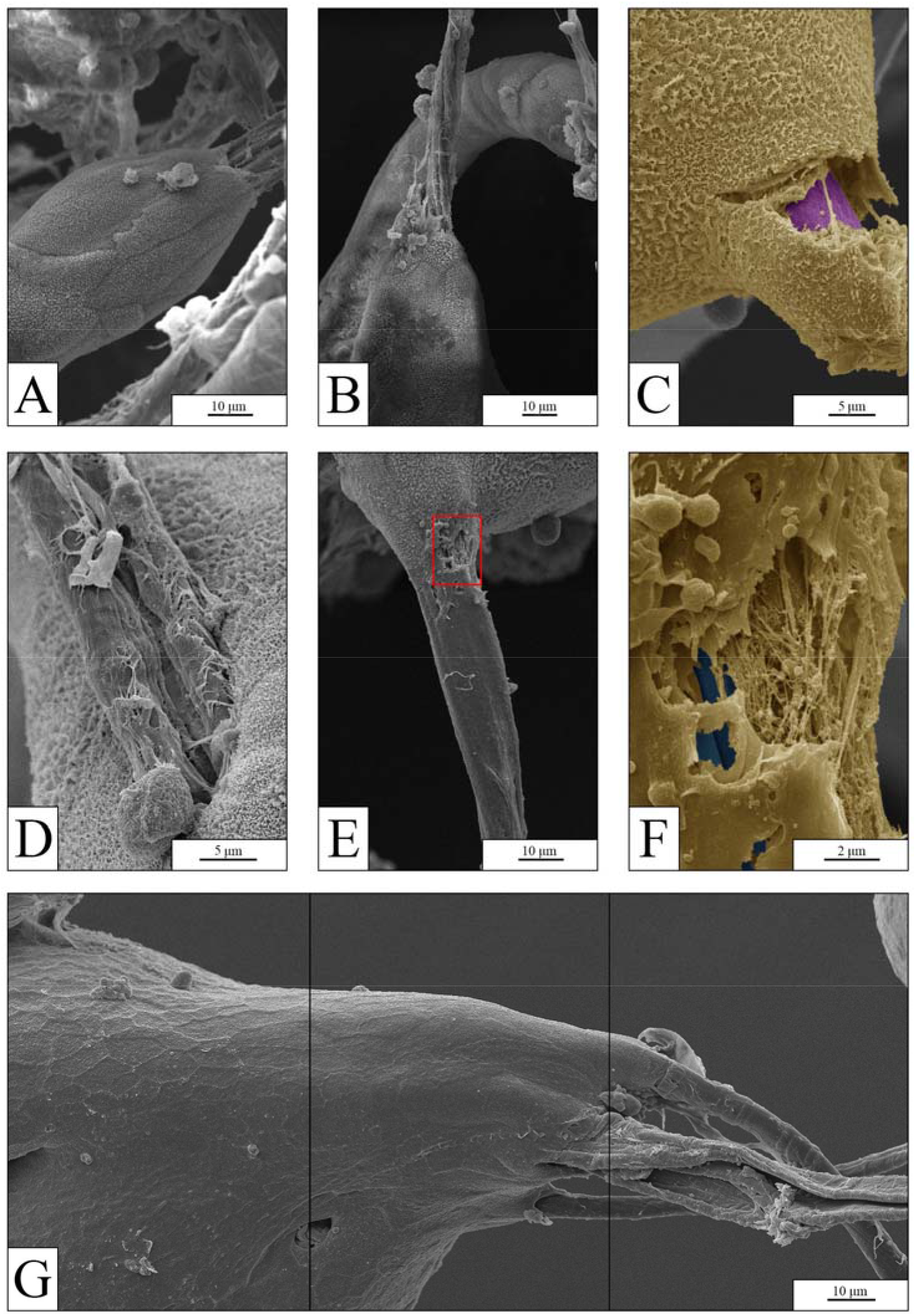
SEM images of human lung organoids exposed to MPFs, partially pseudo-colored for better visualization (cell tissue, yellow; fibers, purple/blue). (A–B) Oriented cell growth of organoids along the length of a fiber; the cell structure and partially coated fibers are shown (MPF concentration: 10 mg L^−1^). (C) Complete internalization of a fiber inside an organoid (MPF concentration: 50 mg L^−1^). (D) Net-like extension of cells growing on a fiber. (E and its magnification F) Thin cell extension covering the surface of a fiber exiting the organoid body (MPF concentration: 50 mg L^−1^). (G) Merged image of three continuous images showing the polarized growth of organoids with the length of multiple fibers (MPF concentration: 50 mg L^−1^).

### Inflammation and oxidative stress evaluation

To investigate possible changes in the expression of inflammatory cytokines and oxidative stress-related genes, gene expression analysis of hAOs was performed at the end of their coculture with the highest MPF concentration. The results revealed no significant differences in the gene expression of cytokines or oxidative stress-related genes in hAOs exposed to MPFs in relation to the control hAOs (Figure 5C-D). In contrast, an increase of inflammatory markers was observed after poly(I:C) treatment, as reported by others (Jose et al. 2020) (Figure 5C).

## Discussion

The first objective of the present study was to set up an innovative *in vitro* model based on the use of 3D structures that could adequately represent the native lung, where emerging atmospheric contaminants such as synthetic materials could have adverse human health effects. And secondly, we wanted to use this model to investigate the potential effects of polyester fibers isolated from a dryer machine on human cell biology.

Organoids have been recently exploited to overcome the limitations of 2D cell culture systems. Due to their capacity to self-organize into minimal biological units and thus potentially recapitulate the functionality and complexity of the tissue of origin, they have emerged as powerful models for studying human development and disease (Fatehullah et al., 2016). Although lung organoids, including those presented herein, do not yet recapitulate all of the complex structures and cellular interactions associated with the different regions of the lung (e.g., trachea, bronchi, bronchioles, alveoli and all mesenchymal and vascular compartments), they contain specific cell types present in the epithelium of the original organ. Therefore, they can be applied for basic and translational research (Barkauskas et al. 2017). We showed that lung-specific cell types, including basal, AT2, and ciliated cells, are represented in our human lung organoids (Figure 7A).

This study presents an attempt to use human lung organoids for toxicological assessment of the effects of MPFs. Synthetic textiles have been identified and extensively described as one of the major sources of MPFs in wastewater (Browne et al., 2011; De Falco et al., 2019). The current research is the first to report the amount and features of MPFs released from synthetic fabrics in a dryer machine, addressing potential atmospheric contamination. In fact, a recent paper by O’Brien et al. (2020) has shown that the use of dryer machines causes an increase of the MPF concentration in the ambient air; however, they did not measure the total amount of MPFs released by the drying process. Washing and drying textiles in household machines present two possible MPF emission pathways: one being through the wastewater (residual water in textiles after the centrifugation cycle of the washing machine), and one being via the air passing by an integrated air filter. The former goes directly into domestic wastewater without filters. Here, we focused on the air pathway; MPFs released by textiles and forced by a strong air flux into the air exhaust are retained by the air filter and then specifically collected at the end of each drying cycle. Considering the evidence reported by O’Brien et al. (2020), the air filter does not remove all MPFs coming from the dryer machine. Thus, the total amount of MPFs released in the air flux can be even higher.

The effects of MPs/NPs on human health have been tested previously using a variety of 2D cell lines. The observed effects were very diverse and depended on the properties of the particle (e.g., size and polymer), the duration and concentration of exposure, and the tested cell line. Interestingly, most of the existing studies analyzing the cytotoxicity of virgin NPs have demonstrated a lack of adverse effects in intestinal, cerebral, and epithelial cell lines. A concentration-dependent significant decrease in cell viability (Lim et al. 2019) and disruption in gene transcription and protein expression after exposure to polystyrene NPs (25 and 70 nm) were only detected for lung cells (Xu et al., 2019), highlighting the need to study the effects of inhaled MPs/NPs. Moreover, with only a few exceptions (Magrì et al. 2018), all previous studies used spherical polystyrene NPs, which are limited in a realistic exposure scenario. In particular, tests representing the human lung should use MPs/NPs in a size and shape that have been shown to be present in the air. Since a substantial portion of airborne environmental MPs consist mostly of MPFs between 200 and 600 μm (Dris et al. 2016; Henry et al. 2019) and fibers >250 μm have been found in human lungs (Pauly et al. 1998), the MPFs applied in the current study represent an environmentally relevant and realistic test material.

Based on a review of studies regarding atmospheric MPs, Prata et al. (2018) calculated that a person’s lungs could be exposed to 26-130 airborne MPs a day. In addition, Vianello et al. (2019) have shown that half of the inhaled and identified MPs are smaller than 5 μm. It has already been reported that inhalability, deposition, and persistence of MPs in the lungs depend on their properties and the different regions of the respiratory tract (Prata 2018). For fibers, in general, deposition in the respiratory tract is well summarized by Mossman et al. (2011). For example, it has been assumed that the length of MPs is of secondary importance because the fibers can align themselves in parallel with the respiratory flow axis and can penetrate to the periphery of the lung and into the alveoli, as already observed in lung biopsies by Pauly et al. (1998). Once they enter deep into the lung, particles (especially the longer fibers) tend to avoid clearance since the magnitude of retention in the lungs depends on the fiber diameter and length (Timbrell 1982). If not cleared by biological mechanisms, MPFs persist in the lung.

The existing database of fiber toxicity studies (Mossman et al., 2011) suggests that human exposure to respirable fibers may induce significant and persistent pulmonary inflammation, cell proliferation, or fibrosis. Immune cells are not the only source of pro- and anti-inflammatory cytokines in the lung; the epithelial cells themselves are known to produce numerous cytokines directly in response to injury or pathogens (Feng et al. 2020), and they contribute to the impressive innate immunity functions of the lung (Whitsett and Alenghat 2015).

Despite the mild and somewhat controversial effects obtained in 2D systems exposed to MPs/NPs, we did not find striking results. Indeed, our findings did not show any significant activation of the inflammatory or oxidative stress pathway at a molecular level. Similarly, confocal microscopy and SEM images did not provide any evidence of organoid growth inhibition or cellular differentiation; these findings were the same as those of the control group. This observation was not completely surprising, as the field of biomaterials is increasingly demonstrating how plastic can be applied as an inert scaffold to support cell growth for human tissue engineering without altering the biological features (Jubeli et al. 2019). Although the biocompatibility of several plastic materials has already been demonstrated, polyester fibers can contain additives that can be added to textiles in order to impart desirable physical, chemical, and biological (i.e., antimicrobial) properties (Sait et al. 2021). Since additives are typically not covalently bonded to the polymer matrix and can therefore leach, they are considered more harmful than the polymer itself, inducing several effects such as endocrine disruption (Kitamura et al., 2005), genotoxicity (Zhao et al., 2013), immunotoxicity (Qiu et al., 2018), and developmental toxicity (Mu et al., 2018). Our results did not show inflammation- or oxidative stress-induced effects of polyester MPFs on hAO development, suggesting that additives, whenever they were present, did not cause adverse effects. On the other hand, the complementary information provided by van Dijk et al. (2021) indicates that for nylon fibers, negative impacts on hAOs were attributed to the potential leaching of nylon components, particularily for organoids under development.

Of note, the most relevant result related to MPF exposure was the polarization of the cell growth along the fibers such that the organoid was induced to envelop the encountered fibers by a cellular layer. This behavior was clearly observed in different cases and at different MPF concentrations, indicating that it should be carefully considered as a possible adverse effect of MPFs. This effect is particularly alarming if we consider repair processes that could follow MPF inhalation, as it is known that fibers can be retained in the lungs. This conclusion goes in line with the complementary observations by van Dijk et al. (2021) who suggested that airborne MPs may be most harmful to young children with developing airways and to people undergoing high levels of epithelial repair.

Based on the findings of this study, during the repair phase of a damaged lung epithelium, the presence of nonbiodegradable fibers may lead to their inclusion in the repaired tissue with unknown long-term effects. This scenario is clearly only a speculation based on an experimental observation performed on a surrogate of the real organ, but it could be the basis for further studies using this promising tool.

## Competing financial interest

The authors declare they have no actual or potential competing financial interests.

## Notes

### Competing Interest Statement

The authors have declared no competing interest.

### Summary of Updates

State of the art in section Introduction and Discussion updated

