## Supplementary figures and images for "Lung organoids and microplastic fibers: a new exposure model for emerging contaminants"

### Supplemental Figure 1

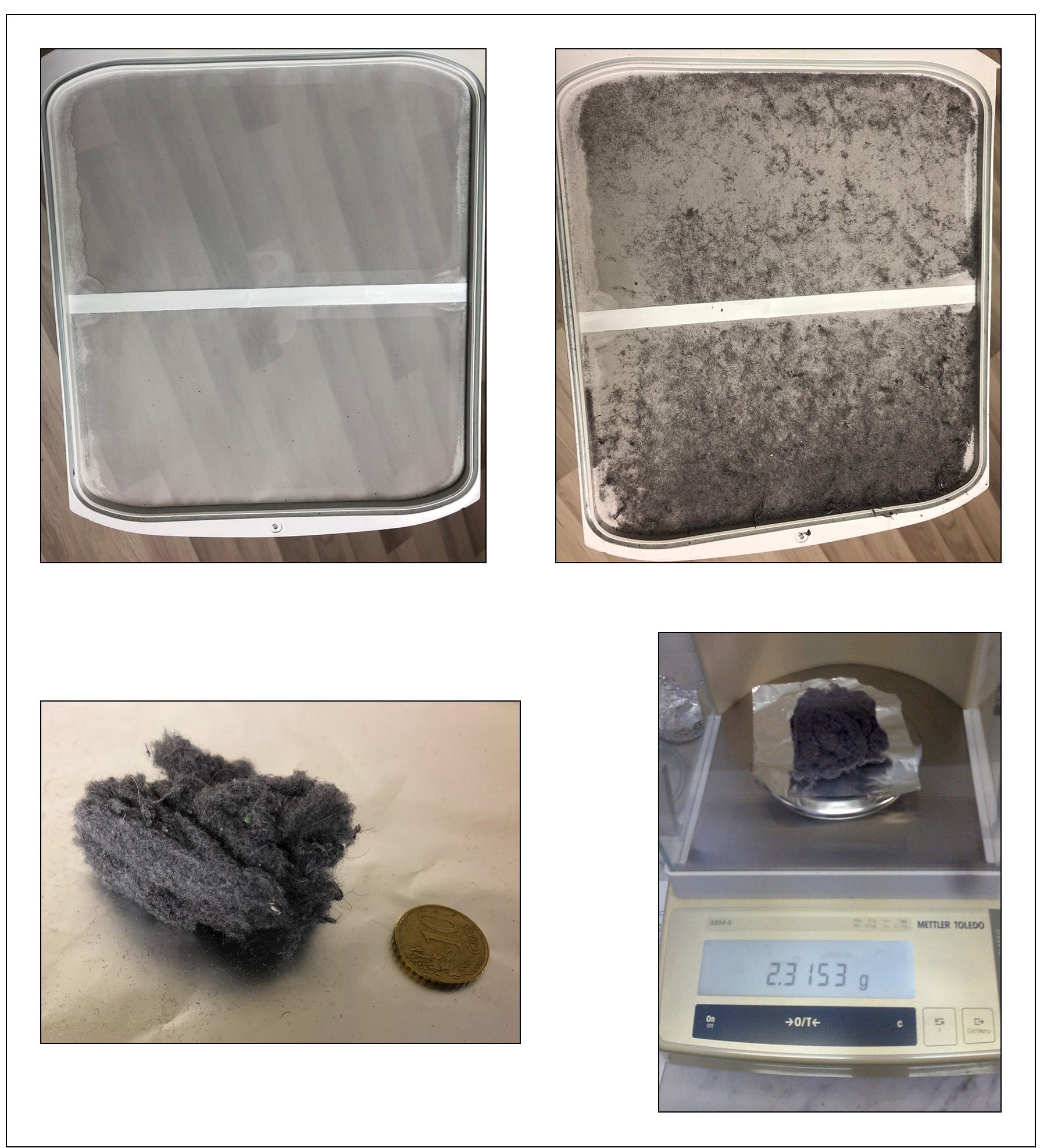

### Supplemental Figure 2

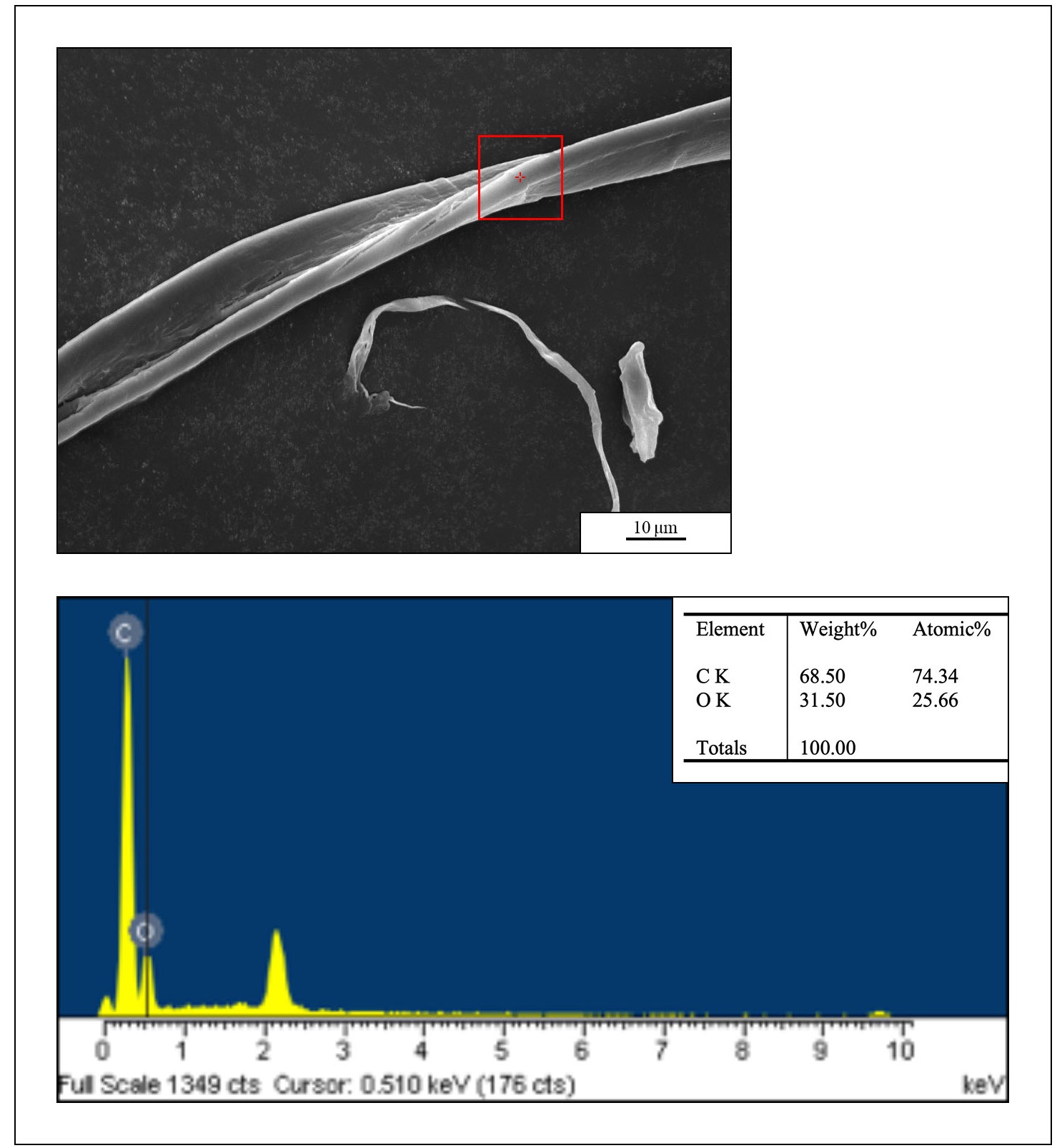

### Supplemental Video 1

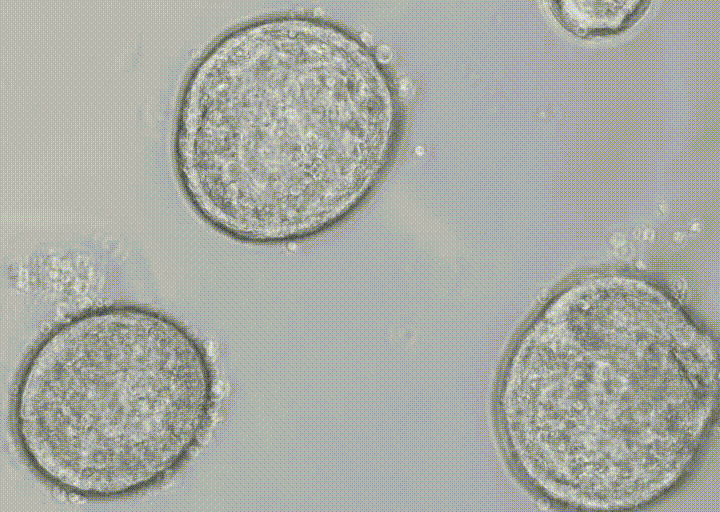
